# Olaparib and Enzalutamide synergistically suppress HCC progression via the AR-mediated miR-146a-5p/BRCA1 signaling

**DOI:** 10.1101/712315

**Authors:** Jie Zhao, Yin Sun, Hui Lin, Fuju Chou, Yao Xiao, Ren’an Jin, Xiujun Cai, Chawnshang Chang

## Abstract

Hepatocellular carcinoma (HCC) is one of most common malignant tumors worldwide, however, the treatment for advanced HCC remains unsatisfactory. Here we found that Olaparib, a FDA approved PARP inhibitor, could enhance the cytotoxicity in HCC cells with a lower BRCA1 expression, and suppressing the AR with either Enz or AR-shRNA could further increase the Olaparib sensitivity to better suppress the HCC cell growth *via* a synergistic mechanism that may involve suppressing the expression of BRCA1 and other DNA damage response (DDR) genes. Mechanism studies revealed that Enz/AR signaling might transcriptional regulting the miR-146a-5p expression *via* binding to the AREs on its 5’ promoter region, which could then lead to suppress the homologous recombination-related BRCA1 expression *via* direct binding to its 3’ UTR of mRNA. Preclinical study using an *in vivo* mouse model also proved that combined Enz plus Olaparib led to better suppression of the HCC progression. Together, these *in vitro/in vivo* data suggest that combining Enz and Olaparib may help in the development of a novel therapy to better suppress the HCC progression.

## Introduction

Hepatocellular carcinoma (HCC) is the most common form of primary liver cancers worldwide. It ranks the 2nd and 6th cause of cancer-related deaths in men and women, respectively^1^. Although HCC at an early stage can be treated effectively through surgery, the prognosis of advanced HCC is still very poor, and more effective treatments for HCC are urgently needed.

HCC frequently arises from chronic inflammation that produces DNA damage^2^. Emerging studies indicated that HCC development and progression can be associated with mis-regulation of DNA damage response (DDR) signaling^3–5^, which may also contribute to HCC’s resistance to ionizing radiation and chemotherapy^5,6^. These evidences suggest that targeting DDR-related signals can be a potential therapeutic approach to suppress HCC progression.

Olaparib is a poly (ADP-ribose) polymerase (PARP) inhibitor that can suppress this key enzyme involved in DDR^7^. In 2014, Lynparza (Olaparib) from AstraZeneca was approved by FDA for patients with deleterious germline BRCA1-mutated advanced ovarian cancer after three or more courses of chemotherapy. In prostate cancer, a phase-2 trial in patients with metastatic lesions also indicated that Olaparib might have a high response rate in patients who had defects in the DDR-related genes, including BRCA2 and ATM^8^. Furthermore, the Phase-3 trial of Olaparib also showed that it might have benefits in patients with germline BRCA mutation and HER2-negative metastatic breast cancer^9^.

Early studies indicated that suppression of PARP1/2, one of the DDR-related genes involved in the base excision repair (BER) pathway for spontaneous single-strand breaks (SSB), might lead to accumulation of the unrepaired SSB, which then results in the collapse of replication forks during S phase and the consequent double-strand breaks (DSB). Normally, these DSBs will be repaired by the homologous recombination repair (HR) machinery. As BRCA1/2 plays a critical role in HR signaling, loss of BRCA function will induce HR deficiency and lead to irreparable DNA damage and cell death in patients when treated with Olaparib.

Enzalutamide (Enz) is a FDA-approved antiandrogen used to treat the metastatic castration-resistant prostate cancer (CRPC)^10,11^. Unlike other antiandrogens, there has been no evidence of hepatotoxicity or elevated liver enzymes in association with Enz treatment in clinical trials^12^.

The androgen receptor (AR) signaling plays important roles in HCC development and progression. Early epidemiological studies indicated that HCC has a gender difference with male to female at 9:2 ratio^13^, and preclinical studies using an *in vivo* HBV-induced HCC mouse model indicated that AR might play positive roles to promote the HBV-induced HCC^14–16^, suggesting that AR might be a potential therapeutic target for HCC^15,16^. Interestingly, recent studies further suggested that AR signaling might regulate the DDR signaling in the various cancer cells^17–19^. Although therapy with antiandrogens alone failed to exhibit therapeutic effect for HCC in early clinical studies^20–22^, those failures suggest the efficacy of antiandrogen therapy might be enhanced if combined with other drugs that can target additional signals.

Here we found that Olaparib might have a better suppression efficacy in those HCC cells with a low BRCA and/or AR expression. Enz altered the BRCA1-related HR signaling, and combining Enz with Olaparib led to better suppress HCC progression in a synergetic manner. Mechanism dissection revealed that Enz/AR might function *via* altering the miR-146a-5p expression to modulate the BRCA1 expression.

## Materials and Methods

### Cell culture

HCC SK-HEP-1 and HepG2 cell lines were obtained from ATCC and authenticated by a professional Biotechnology Company in 2015. The HA22T cell line (BCRC No. 60168) was a gift from Prof. Yuh-Shan Jou, Academia Sinica, Taiwan^41^. The LM3 cell line was obtained from the Type Culture Collection of the Chinese Academy of Sciences (Shanghai, China). The HCC cells were cultured in Dulbecco’s Modified Eagle’s Medium (Invitrogen) with 10% FBS, 1% glutamine, and 1% penicillin/streptomycin. Cells were cultured in a 5% (v/v) CO_2_ humidified incubator at 37°C.

### Plasmids and Lentivirus

PLVTHM, pLKO.1, pWPI-GFP lentivirus vectors were used to overexpress miR-146a-5p and introduce short hairpin RNAs (shRNAs), for knocking down AR and overexpressing AR cDNA, respectively. To generate stable transfectants, the lentivirus vector, and pMD2G envelope plasmid, the psAX2 packaging plasmid, were transfected into 293HEK cells using the standard calcium chloride transfection method. The lentivirus soup was collected after incubating 48 and 72h, concentrated and used immediately or frozen in −80°C for lateruse.

### MTT cell viability assay

HCC cells were placed in 24-well plates at a density from 2×10^3^ to 5×10^3^ cells/well. After drug treatment, culture media was removed and 500 μl MTT (0.5 mg/mL) per well was added and incubated for 1 hour. The absorbance at 570nm was detected. Cell viability was calculated using the formula, [OD(sample)-OD(blank)]/[OD(control)-OD(blank)]. At least triplicate experiments were performed, and values with mean ± S.D. were presented.

### Western blot

Cells were lysed in ice-cold cell lysis buffer and equal amounts of the protein were loaded for electrophoresis on denaturing SDS/PAGE gel and then transferred onto PVDF membranes (Millipore, Billerica, MA). The blots were probed with the primary antibodies overnight at 4 °C, followed by incubation with the appropriate secondary antibodies at room temperature for 1 h. The bands were visualized using an ECL chemiluminescent detection system (Thermo Fisher Scientific, Rochester, NY).

### Quantitative real-time PCR (qRT-PCR)

For mRNA detection, total RNAs were isolated using Trizol reagent (Invitrogen). One μg of total RNA was subjected to reverse transcription using Superscript III transcriptase (Invitrogen). The qRT-PCR was conducted using a Bio-Rad CFX96 system with SYBR green to determine the expression level. Expression levels were normalized to GAPDH level. Primers were listed in Supplementary Table.

For miRNA, briefly, 500 ng RNAs were processed for poly A addition by adding 1 unit of polymerase with 1 mM ATP in 1xRT buffer at 37 °C for 10 min in 10 μl volume, then heat inactivated at 95°C for 2 min, add 50 pmol anchor primer to 12.5 μl, and incubated at 65°C for 5 min. For cDNA synthesis, we added 2 μl 5x RT buffer, 2 μl 10 mM dNTP, 1 μl reverse transcriptase to total 20 μl, and incubated at 42 °C for 1 h. The miRNA levels were detected by TaqMan assay. Expression levels were normalized to the level of U6.

### RNA Immunoprecipitation (RIP)

Total RNA was isolated using Trizol reagent (Invitrogen) and dissolved in RIP buffer (150 mM KCl, 25 mM Tris/pH 7.4, 5 mM EDTA, 0.5 mM DTT, and0.5% NP40) with RNase inhibitor and protease inhibitor cocktail. The RNA was cleared by protein A/G beads for 1 hour and incubated with Argonaute-2 antibody or IgG for 4 hours at 4 °C. Then the beads pre-blocked with 15 mg/ml BSA were added to the antibody-lysate mixture and incubated for another 2 hours. The RNA/antibody complex was washed 3 times by RIP buffer. The RNA was extracted using Trizol (Invitrogen) according to the manufacturer’s protocol and subjected to qRT-PCR analysis.

### Luciferase reporter assay

The wild type and mutant 3’ UTR of BRCA1 were constructed into psi-check-2 vector (Promega, Madison, WI, USA). Cells were plated in 24-well plates and the plasmids (psi-check-2-WT, psi-check-2-mut) were transfected using Lipofectamine 3000 (Invitrogen) according to the manufacturer’s instructions. Luciferase activity was measured by Dual-Luciferase Assay reagent (Promega) according to the manufacturer’s manual.

The wild type and mutant promoter of miR-146a-5p were constructed into pGL3 vector (Promega). The pGL3-WT or pGL3-mut were co-transfected with pRL-TK using lipofectamine^®^ 3000. Luciferase activity was measured by Dual-Luciferase Assay reagent (Promega) according to the manufacturer’s manual.

### Chromatin Immunoprecipitation Assay (ChIP)

After cross-linking with formaldehyde, the cells were lysed, the fraction containing nuclear pellets was isolated and chromatin was sonicated to an average fragment size of 200–1000bp. Samples were pre-cleared with protein A-agarose. The precleared samples were incubated with AR antibody or IgG for 4 hours at 4 °C. Then the pre-blocked beads were added to the antibody-lysate mixture and incubated for another 2 hours. After several washes and de-crosslinking, the DNA was collected using QIAprep spin miniprep columns. Specific primer sets were designed to amplify a target sequence within human miR-146a-5p promoter.

### Alkaline comet assay

Cells were grown, treated, and irradiated in the described conditions. Damaged DNA was detected using alkaline CometAssay^®^ (TREVIGEN, Gaithersburg, MD), according to the manufacturer’s instructions. Slides were stained with SYBR Gold (S11494, Thermo Fisher Scientific) and visualized using a fluorescence microscope. The comet tail was scored according to DNA content. Multiple randomly selected cells were analyzed per sample.

### Homologous Recombination (HR) DR-GFP Assay

1 × 10^6^ SK-HEP-1 cells in 6-well plates were pre-treated as described. Cells were transfected with 0.5 μg of pDR-GFP plasmid (Addgene #26475) and 2 μg of I-SceI expression plasmid pCBASceI (Addgene #26477) using lipofectamine® 3000 (#11668019, ThermoFisher Scientific). After 48 hrs, GFP positive and total cells were counted in 3 random high power fields under a fluorescence microscope.

### Animal experiment

The animal experiment was based on the Zhejiang University guide for the care and use of laboratory animals. 5 × 10^6^ SK-HEP-1-luc cells were implanted into subcutaneous pockets in the flanks of 6 week-old male nude mice. After 6 weeks, the tumors were removed and cut into pieces. Equal-sized tumor pieces were implanted into livers of other 6 week-old male nude mice. After 4 weeks, the orthotopic tumor was detected by bioluminescent images. The nude mice were then randomly divided into four treatment groups: vehicle, Enz, OLA and Enz+OLA groups (Enz, 10 mg/kg per day, orally; OLA, 40 mg/kg per day, intraperitoneally). Each group had three mice. The orthotopic tumor growth was monitored weekly by bioluminescent images. Mice were sacrificed after 3 weeks treatment. Tumor specimens were collected for further analysis.

### Immunohistochemical analysis

Tumor specimens were fixed in formalin, embedded in paraffin, and sectioned at a thickness of 4 mm for immunohistochemical analysis of BRCA1, ki67 and cleaved caspase-3. Rehydrated slides were microwave-heated for 20 min in citrate buffer (10 mM/pH 6.0) for antigen retrieval and then incubated with 1% H_2_O_2_ for 10 min to inactivate endogenous horseradish peroxidase (HRP). After blocking with serum-free protein block (Dako), they were incubated with the primary antibodies for 120 min at room temperature, followed by incubation with HRP-conjugated secondary antibody (Dako) for 60 min at room temperature. Chromogenic reactions were carried out according to the protocols of the ImmPACT™ DAB Kit (Vector Laboratories). For quantitative analyses, 6 microscopic fields (at 200×) of tumors were randomly selected. The labeling rates of BRCA1, Ki67 and cleaved caspase-3 were evaluated.

### Statistical Analysis

Experiments were performed in triplicate and reapeated at least 3 times. Data are presented as mean ± SD or SEM from independent experiments. Statistical analyses involved linear regression analysis, Student’s t test, one-way ANOVA tests, two-way ANOVA tests and Log-rank (Mantel-Cox) test with SPSS 22 (IBM Corp., Armonk, NY) or GraphPad Prism 6 (GraphPad Software, Inc., La Jolla, CA). P < 0.05 was considered statistically significant.

## Results

### Differential sensitivity of HCC cells to Olaparib treatment

We first applied the MTT assay to examine the cytotoxic effect of Olaparib in multiple human HCC cell lines, including HA22T, SK-HEP-1, HepG2, and LM3 cells. The results revealed that treating with 5 μM Olaparib for 3 days resulted in different cell viabilities with a differing relative OD value of Olaparib compared to vehicle control, showing HepG2 (OD=68.2%) and LM3 (OD=69.6%) were more sensitive to Olaparib than HA22T (OD=88.9%) and SK-HEP-1 (OD=83.6%) cells, P= 0.0067 **(Fig. 1A)**.

**Fig 1.**
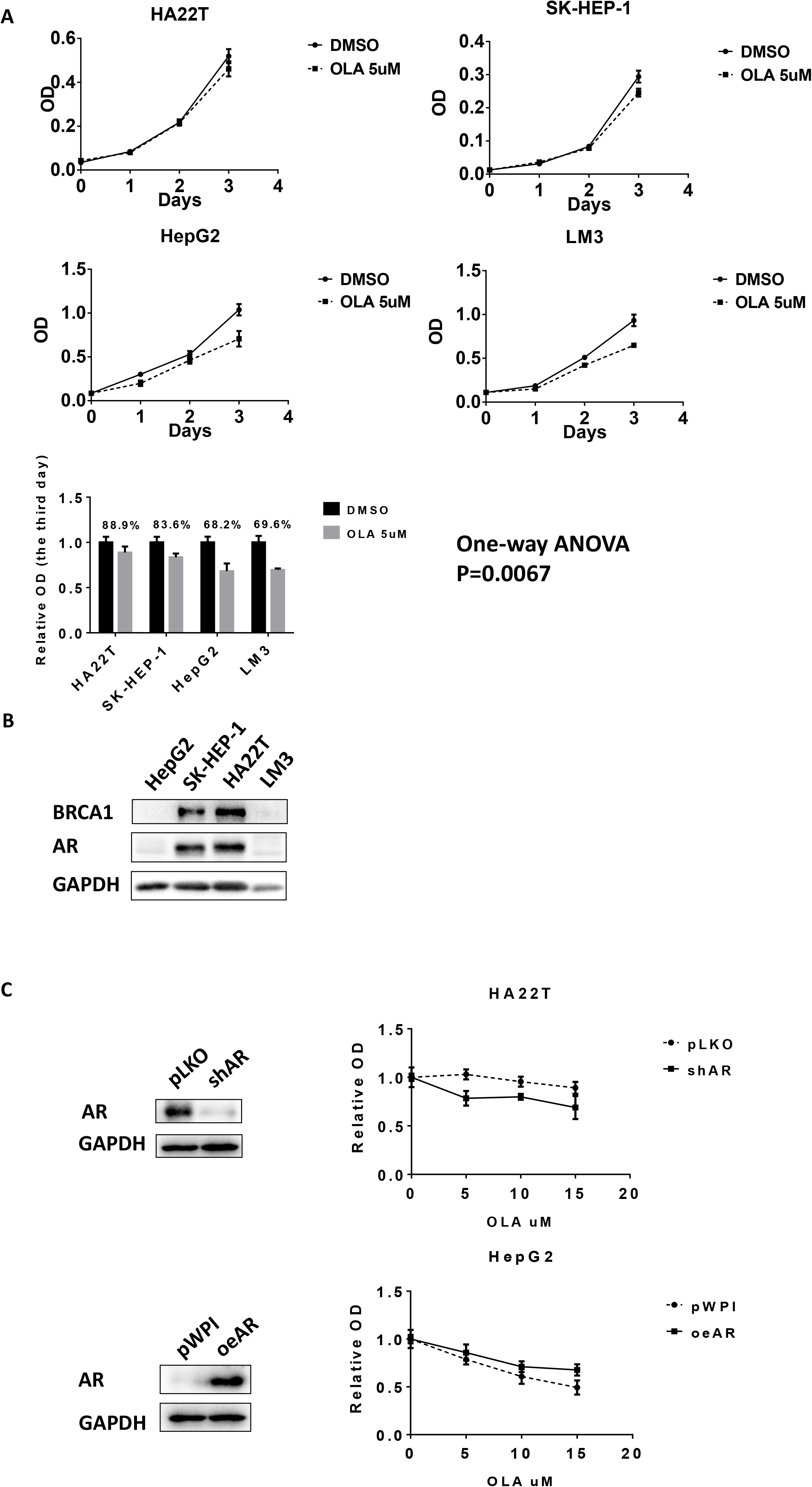
Differential sensitivity of HCC cells to Olaparib treatment. (A) Human HCC cell lines HA22T, SK-HEP-1, HepG2, and LM3 were treated with DMSO or 5μM Olaparib. Cell viability was detected using MTT assay with quantitation in lower panel. (B) Western blots showed protein expression levels of BRCA1 and AR in different HCC cell lines. (C) HA22T and HepG2 cells were stably infected with lentiviruses expressing shRNA targeting AR or AR cDNA for overexpression AR. Western blots (left) showed the expression of AR and GAPDH (loading control). After manipulating AR, cells were treated with different concentrations of Olaparib for 2 days. The MTT assay was performed to determine cell viability (right).

To further study why Olaparib has different sensitivity in different HCC cells, we focused on the BRCA1 expression since recent studies indicated that BRCA-defective cancer cells are more sensitive to Olaparib^23^. Using western blot analysis, we found Olaparib-less-sensitive HCC HA22T and SK-HEP-1 cells are “BRCA1-positive”. In contrast, Olaparib-more-sensitive HepG2 and LM3 cells are “BRCA1-negative” **(Fig. 1B)**.

As recent findings indicated that the AR could induce “BRCAness” in prostate cancer (PCa), we also determined AR expression in different HCC cell lines. Interestingly, we found that AR expression was also associated with BRCA1 expression in different HCC cell lines, showing “AR-negativity” is also related to “BRCAness” **(Fig. 1B)**.

To test whether AR signaling is crucial for Olaparib sensitivity in HCC, we further manipulated AR and tested the Olaparib sensitivity of the HCC cells through MTT assay. The results revealed that suppressing AR in HA22T cells increased the Olaparib sensitivity and increasing AR in HepG2 cells decreased the Olaparib sensitivity in the HCC cells **(Fig. 1C)**.

Together, results from **Fig. 1A-C** suggest that AR may play key role to alter the Olaparib sensitivity in HCC cells that may involve the linkage of AR and BRCA1.

### Mechanism dissection of how AR can influence the Olaparib sensitivity: via altering the DDR and HR signals

To dissect the mechanism of how AR may link the BRCA signaling to influence the Olaparib sensitivity of the HCC cells, we applied the human clinical survey *via the* TCGA database analysis, and found a positive correlation between AR *vs* BRCA2 and RAD51, two HR-related proteins^24^, in the HCC patients **(Fig. 2A)**, suggesting AR might function *via* altering the HR signaling including the BRCA1/2 to influence Olaparib sensitivity of HCC cells.

**Fig 2.**
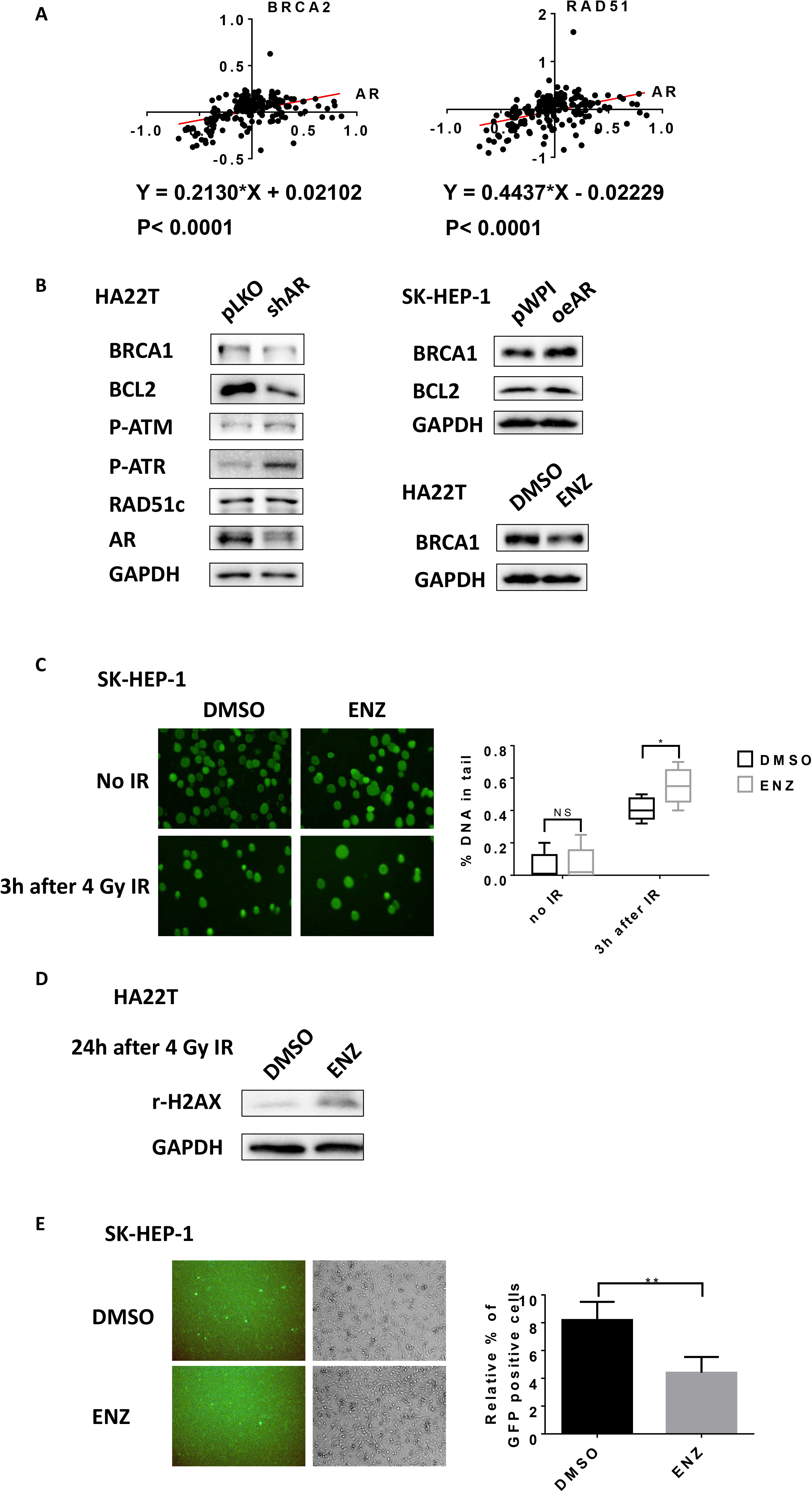
Antiandrogen Enz suppresses BRCA1 and HR pathway. (A) The protein expression correlation between AR and BRCA1, RAD51 in 184 HCC patients based on TCGA database. (B) HA22T cells were infected with lentivirus pLKO control or shRNA targeting AR and Western blots showed some key HR proteins expression levels (left). SK-HEP-1 cells were infected with lentiviruses pWPI control or oeAR. Western blots showed proteins expression levels of BRCA1 and BCL2 (right upper). HA22T cells were treated with 10 μM Enz (ENZ), western blots showed proteins expression levels of BRCA1 (right lower). (C) SK-HEP-1 cells were treated with DMSO or 20 μM ENZ for 24 hours, then irradiated by 4 Gy IR. Three hours after radiation, DNA damage was detected by Alkaline CometAssay^®^. The comet tails were scored according to DNA content. Multiple randomly selected cells were analyzed per sample. (D) HA22T cells were pretreated with DMSO or 20μM ENZ for 24h, followed by 4 Gy IR. 24 hours after radiation, r-H2AX and GAPDH were detected by western blot. (E) SK-HEP-1 cells were pretreated with DMSO or 20 μM Enzalutamide for 24 hours and were co-transfected with pDRGFP and I-SceI plasmids. After 48 hours, the GFP+ cells were observed under fluorescence microscope. The proportion of GFP+ cells were presented as relative percentage. Five randomly selected microscopic fields were analyzed per sample. Quantitation data are presented as mean ± SD. *P < 0.05, **P < 0.01, NS=Not Significant.

These human clinical results were further confirmed in the *in vitro* studies, showing that suppressing AR *via* adding AR-shRNA (shAR) also led to decrease the expression of BRCA1 and BCL2 **(Fig. 2B left)**, two well-known factors that can influence HR and HCC cell survival. In contrast, increasing AR *via* adding AR-cDNA (oeAR) led to increase the expression of BRCA1 and BCL2 **(Fig. 2B right upper).**Similar results were also obtained when we replaced AR-shRNA with 10 μM antiandrogen-Enzalutamide (Enz) **(Fig. 2B right, lower)**.

Using the comet assay^25^ to examine the influence of Enz on the DDR process, we found treating with 20 μM Enz led to more DNA damage 3 hours after ionizing radiation **(Fig. 2C)** in SK-HEP-1 cells. Also, treating HCC HA22T cells with 20 μM Enz 24 hrs after radiation-treatment resulted in accumulation of much more r-H2AX **(Fig. 2D)**.

Importantly, using the HR DR-GFP assay^26^ to specifically analyze the influence of Enz on the HR, we found that treating the SK-HEP-1 cells with 20 μM Enz led to decrease the GFP positive cells, therefore a reduction in successful DDR through HR **(Fig. 2E)**.

Together, results from multiple approaches (**Fig. 2A-D)** all conclude that Enz/AR signaling can alter the BRCA-related DDR/HR signals in the HCC cells.

### Combining Enz with Olaparib resulted in better suppression of the HCC cell viability *via* a synergistic effect

Since Olaparib may have better therapeutic effect on those “BRCA1-negative” HCC cells and Enz treatment can effectively suppress BRCA1 expression, we were interested to see if the combined Enz and Olaparib could synergistically suppress the “AR-positive” HCC cell growth. The results revealed that combining Enz and Olaparib led to better suppression of the cell growth in the two HCC cell lines, HA22T and SK-HEP-1, and the CompuSyn software**^27^**analysis confirmed that these two currently clinically used drugs had synergistic cytotoxic effects to suppress the HCC cell growth **(Fig. 3A)**. Similar results were also obtained when we applied another MTT assay showing dramatic reduction of cell viability and proliferation in the HCC SK-HEP-1 cells (Two-way ANOVA, P<0.05) **(Fig. 3B)**.

**Fig 3.**
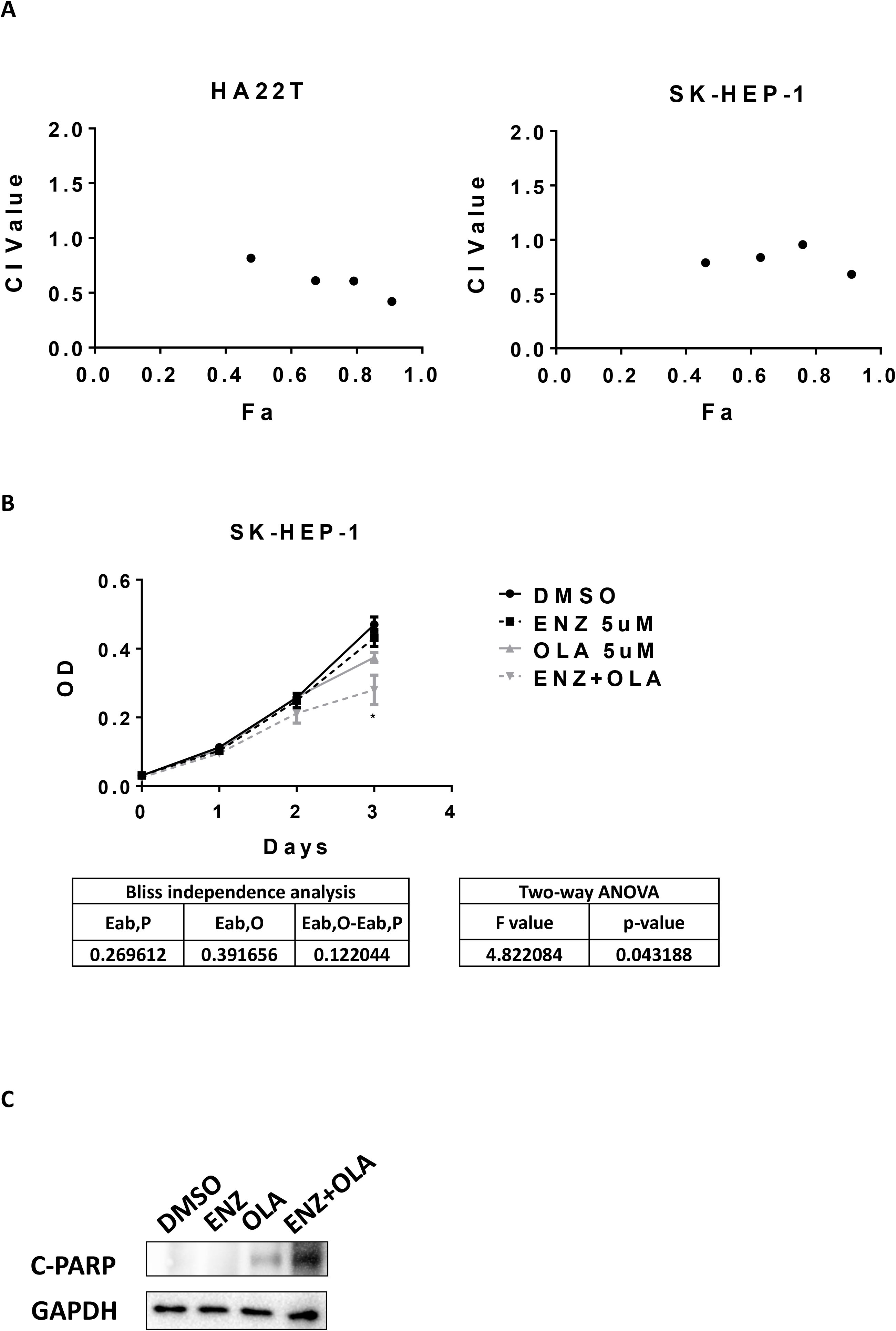
Enz and Olaparib synergistically suppress HCC. (A) HA22T and SK-HEP-1 cells were treated with serial dilutions of Enz (ENZ) and Olaparib (OLA) alone or with combination in the same concentration ratio of 4:3 for 5 days. The MTT assay was performed to determine cell viability. Data was analyzed using CompuSyn software based on median-effect principle and combination index theorem (CI<1, =1, and >1 indicates synergism, additive effect and antagonism, respectively). (B) SK-HEP-1 cells were treated with 5 μM ENZ and 5 μM OLA alone or with combination. The MTT assay was performed to determine cell viability. Synergistic effects were determined by two-way ANOVA tests and by the Bliss independence model. (C) SK-HEP-1 cells were treated with 10 μM ENZ and 10 μM OLA alone or in combination for 48 hours. The protein levels of cleaved PARP (C-PARP) and GAPDH were detected using western blot.

Results from the apoptosis assay to detect the level of cleaved PARP (c-PARP) also revealed that combining Enz with Olaparib could dramatically increase c-PARP level, suggesting that synergistic cytotoxicity of combining Enz with Olaparib treatment could be the result of synergistic apoptosis in HCC cells **(Fig 3C)**.

Together, these multiple *in vitro* results (**Fig. 3A-C**) demonstrate that combining Enz and Olaparib treatment can lead to better suppression of the HCC cell growth *via* a synergistic mechanism.

### Mechanism dissection of how Enz/AR signaling can suppress BRCA1 expression: *via* altering the miR-146a-5p expression in HCC cells

To examine the molecular mechanism underlying AR’s regulation of BRCA1 and BCL2, we began by determining the transcriptional regulation using qRT-PCR, and results revealed that altering AR expression with either adding AR-cDNA or AR-shRNA or Enz treatment had not significant impacts on mRNA expression of BRCA1 and BCL2 **(Fig. S1A)**. In addition, it appeared to be not related to regulation of protein stability using the cycloheximide (CHX) assay **(Fig. S1B)**.

We then focused on the role of non-coding RNAs, such as miRNAs (located in the Argonaute 2 complex), in the regulation of BRCA1 and BCL2. We detected the BRCA1 and BCL2 mRNA expression in Argonaute 2 complex using RIP assay with the same levels of Argonaute 2 being pulled-down during the RIP assay **(Fig. S1C)**, yet suppressing AR in HA22T cells could increase BRCA1 and BCL2 expression while increasing AR in SK-HEP-1 cells resulted in a decrease of BRCA1 mRNA in Argonaute 2 complex **(Fig. 4A)**, suggesting that Enz could reduce BRCA1 and BCL2 likely through altering miRNAs to contribute to the synthetic lethality of the two drugs.

To identify the specific miRNAs involved in AR’s regulation of BRCA1 and BCL2, we analyzed miRNA databases and predicted 11 candidate miRNAs that might target BRCA1 and BCL2, with miR-146a-5p significantly increased after altering the AR expression *via* adding AR-shRNA to HA22T cells, **(Fig. 4B left)**, or adding AR-cDNA (oeAR) to SK-HEP-1 cells or treating HA22T cells with 10 μM Enz **(Fig. 4B right)**.

**Fig 4.**
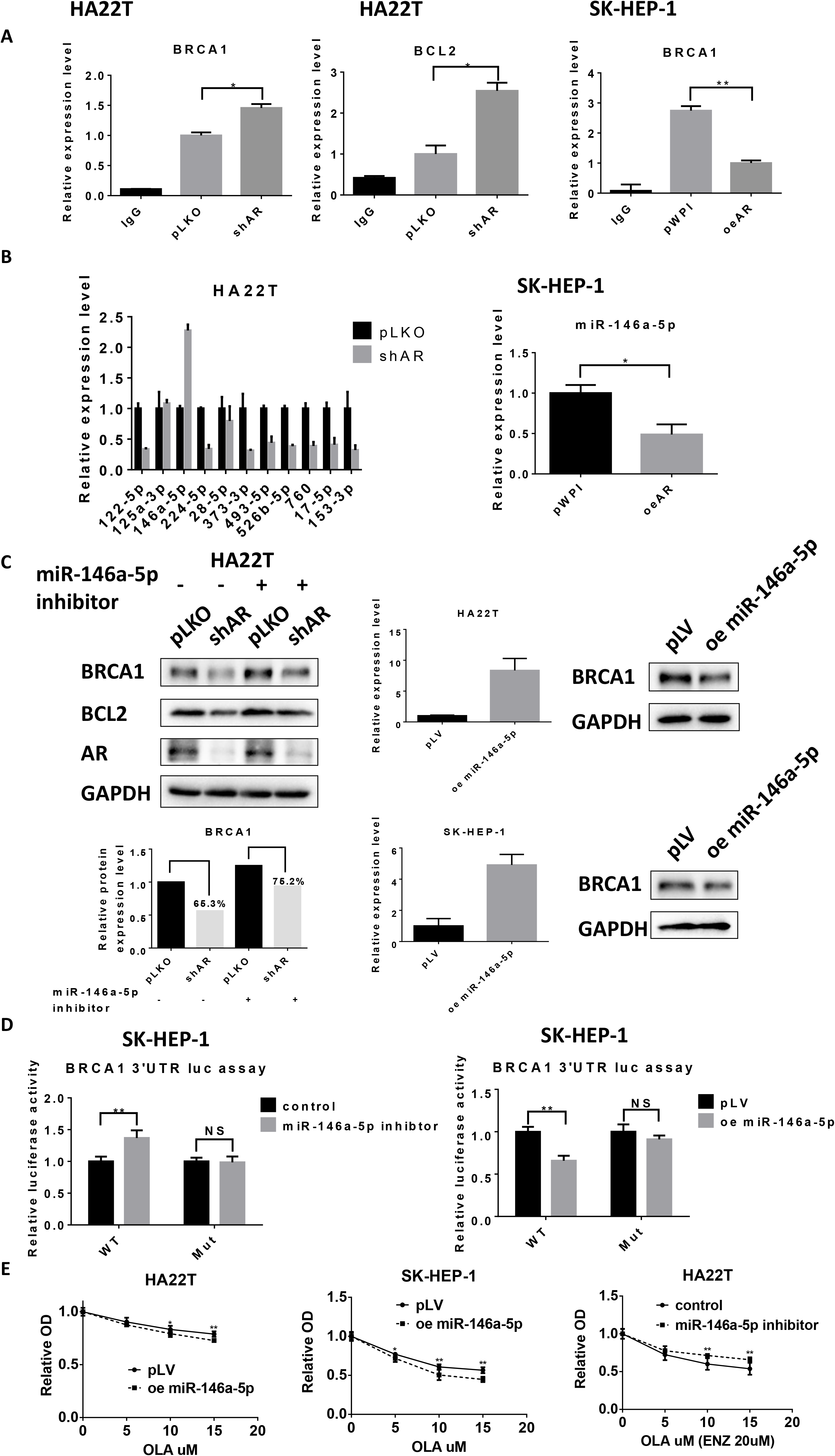
Enz reduces BRCA1 through miR-146a-5p in HCC cells. (A) HA22T cells were infected with lentivirus pLKO control or shAR. SK-HEP-1 cells were infected with lentiviruses pWPI control or oeAR. Anti-Argonaute 2 RNA immunoprecipitation (RIP) was used to detect BRCA1 and BCL2 mRNA expression levels in Argonaute 2 complex by qRT-PCR. (B) In pLKO and AR knocked down HA22T cells, the differential expression levels of 11 candidate miRNAs were detected using qRT-PCR (left). MiR-146a-5p expression level was detected in pWPI, oeAR HepG2 cells (middle). The miR-146a-5p expression levels were detected in HA22T cells treated with DMSO or 10 μM Enz (ENZ) (right). (C) PLKO control or shAR HA22T cells were transfected with miR-146a-5p inhibitor or control, respectively. The protein levels of BRCA1, BCL2, AR and GAPDH were detected using western blot (left upper) and quantified by ImageJ software (left lower). HA22T cells were infected with lentiviruses pLV control or overexpressed miR-146a-5p and expression levels detected using qRT-PCR (middle). BRCA1 and GAPDH protein levels were detected by western blot (right). (D) The wild type (WT) or mutant BRCA1 3’UTR was subcloned into the psiCHECK™-2 Vector. SK-HEP-1 cells were co-transfected with WT or mutant vector, miR-146a-5p inhibitor or control, oemiR-146a-5p or pWPI for 2 days. Luciferase activities were presented in ratios of Renilla to Firefly luciferase reporter activities. **P<0.01, NS=Not Significant. (E) HA22T (left) and SK-HEP-1 (middle) cells transfected with PLV control or oemiR-146a-5p were treated with different concentrations of Olaparib (OLA) for 72 hours. HA22T (right) cells were transfected with miR-146a-5p inhibitor or control and treated with 20 μM Enz (ENZ) and different concentrations of OLA for 72 hours. The MTT assay was performed to determine cell viability. Quantitation data are presented as mean ± SD. **P < 0.01, NS=Not Significant.

Results from a human clinical survey using TCGA databases analysis revealed that miR-146a-5p was an important biomarker in HCC related to tumorigenesis and prognosis. The miR-146a-5p was significantly lower in cancer tissues compared with para-cancerous tissues **(Fig. S1D left)** and patients with a higher miR-146a-5p had better relapse-free survival (RFS) than those with a lower expression **(Fig. S1D right)**.

Using an inhibitor of miR-146a-5p, we confirmed that miR-146a-5p could target BRCA1 and BCL2 **(Fig. 4C left)**. As we are primarily interested in the potential synergistic effect of Olaparib and Enz, we then focused only on BRCA1 for the remainder of the work. We found that the decrease of BRCA1 caused *via* suppressing AR was reversed by miRNA inhibitor **(Fig. 4C left)**, consistent with its role in the regulation of BRCA1. We further overexpressed miR-146a-5p in HA22T and SK-HEP-1 cells and found that overexpression of miR-146a-5p decreased BRCA1 protein level using western blot assay **(Fig. 4C right)**.

### Mechanism dissection of how AR/miR-146a-5p signaling can suppress BRCA1 expression: *via* direct binding to the 3’UTR of BRCA1 mRNA in HCC

Using the psiCheck2 luciferase reporter assay, we further confirmed that miR-146a-5p could target the 3’UTR of BRCA1 mRNA. In the wild type 3’UTR of BRCA1, miR-146a-5p inhibitor increased relative luciferase activity while overexpressing miR-146a-5p decreased it. On the other hand, the mutant 3’UTR of BRCA1 without the predicted binding site failed to respond to miR-146a-5p in either condition **(Fig 4D)**.

In addition, HA22T and SK-HEP-1 cells with overexpressed miR-146a-5p, as expected, became more sensitive to Olaparib **(Fig. 4E left and middle)**. In HA22T cells, the inhibitor for miR-146a-5p decreased cell sensitivity to the combination treatment of Olaparib and Enz **(Fig. 4E right)**.

Together, these data from **Fig. 4A-E** demonstrated that miR-146a-5p could target BRCA1 3’UTR to contribute to the regulation of BRCA1 by AR signaling.

### Mechanism dissection of how AR signaling can suppress miR-146a-5p expression: *via* transcriptional regulation

Next, to determine how AR can regulate miR-146a-5p expression, we hypothesized that AR could directly bind to the promoter of miR-146a-5p to influence its transcription. There are two predicted androgen-response-elements (AREs) within 2 kb of the miR-146a-5p promoter region **(Fig. 5A upper)**. Chromatin immunoprecipitation (ChIP) *in vivo* binding assay with gel format **(Fig. 5B left)**and quantitation format (QPCR quantitation) indicated that AR might bind to the ARE2 instead of ARE1 **(Fig. 5B right)**.

**Fig. 5.**
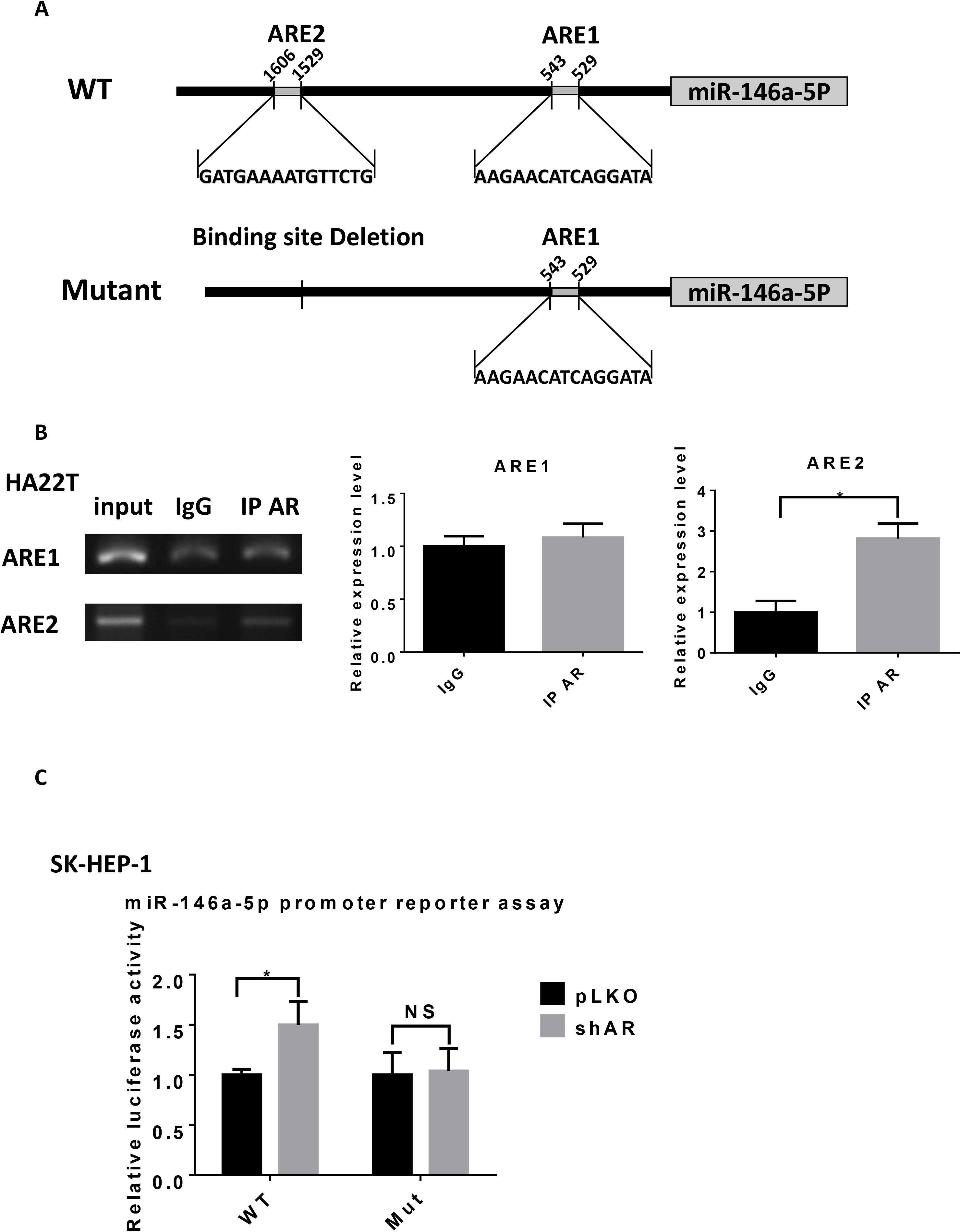
AR directly binds with the promoter of miR-146a-5p to suppress its transcription. (A) Illustration for wild type and mutant promoter regions of miR-146a-5p. (B) In HA22T cells, DNA fragment binding with AR was pulled-down using ChIP assay. Semi-quantitative PCR (left) and qPCR (right) analyses using primers that flanked the AR DNA binding site (ARE1 or ARE2) in the promoter region of the miR-146a-5p. (C) PLKO or shAR SK-HEP-1 cells were transiently co-transfected with luciferase reporter plasmid (pGL3 WT or pGL3 mutant) and pRL-TK vector (control) for 2 days. Luciferase activities were presented in ratios of Firefly to Renilla luciferase reporter activities. Quantitation data are presented as mean ± SD. *P < 0.05, NS=Not Significant.

Using pGL3 luciferase reporter assay, we further confirmed that AR regulated miR-146a-5p through direct binding to the ARE2. In the wild type promoter of miR-146a-5p, knocking down AR increased relative luciferase activity. However, in the mutant promoter without ARE2 **(Fig. 5A lower)**, knocking down AR failed to change the relative luciferase activity **(Fig. 5C)**.

Together, these data from **Fig. 5A-C** suggested that AR signaling may regulate miR-146a-5p *via* direct binding to the ARE2 located on its 5’ promoter region.

### Preclinical study using *in vivo* mouse model to demonstrate combined Enz and Olaparib can lead to better suppression of HCC progression

Finally, we studied the synergistic effect of Enz and Olaparib using the *in vivo* mouse model, and established subcutaneous transplanted SK-HEP-1-luc. HCC cell line model in nude mice. After 6 weeks, tumors were removed and cut into small pieces for orthotopic transplantations in the livers of other nude mice. After 4 weeks, the orthotopic tumor was detected by bioluminescent imaging and the mice were randomly divided into four groups and treated with Enz or Olaparib alone and in combination. The orthotopic tumor growth was monitored weekly by bioluminescent IVIS imaging. After three weeks, we analyzed the data and found that Enz or Olaparib alone could control tumor growth whereas Enz plus Olaparib could significantly shrink tumors **(Fig. 6A)**. Treatment with Enz and Olaparib achieved greater tumor suppression than treatment with either Enz or Olaparib alone. Synergy analysis using Bliss independence model revealed a synergistic therapeutic effect in Enz plus Olaparib group, but P>0.05 when using two-way ANOVA analysis **(Fig. 6B)**.

**Fig 6.**
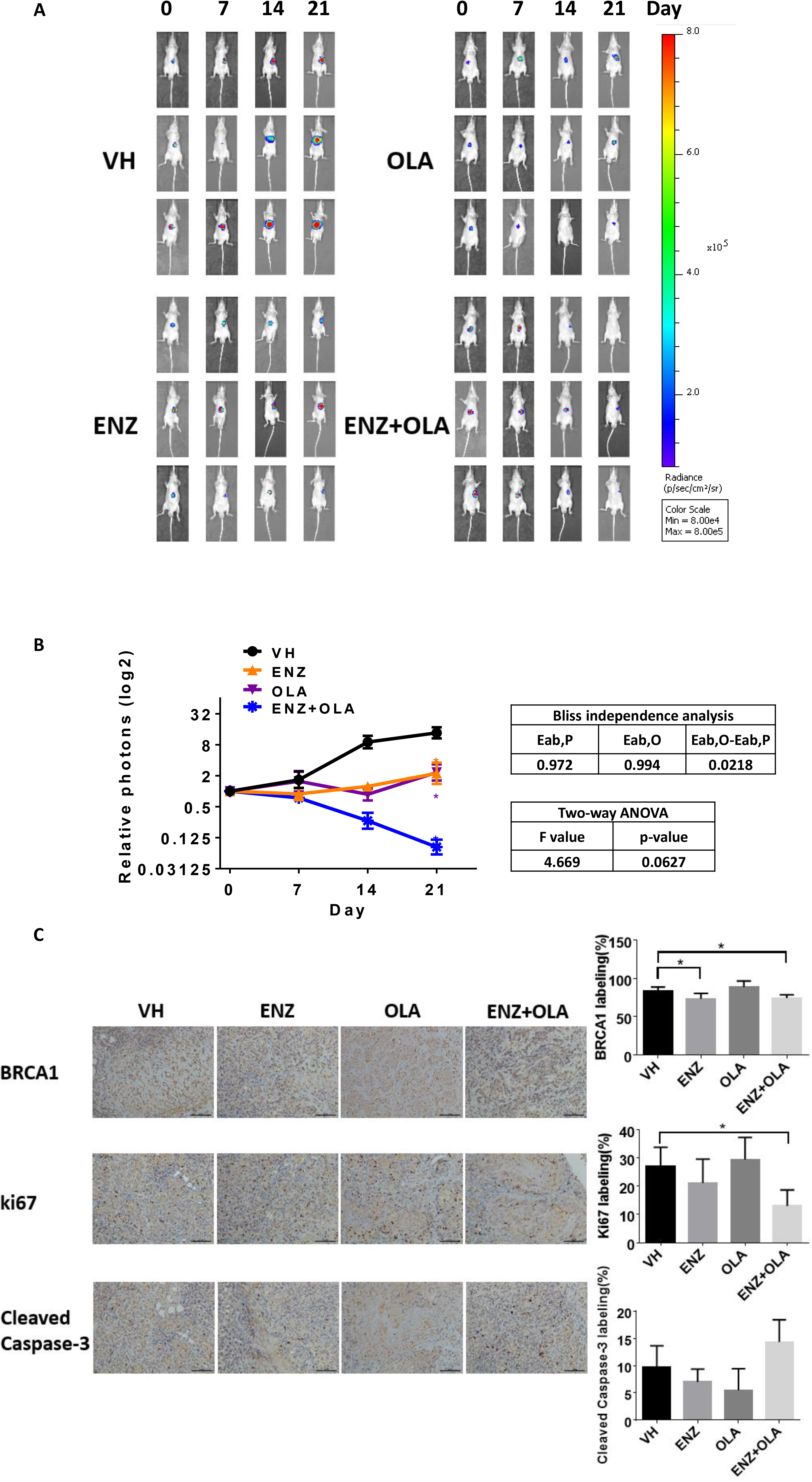
Enz plus Olaparib can lead to better suppression of the HCC growth *in vivo*. (A) SK-HEP-1-luc orthotopic xenotransplanted HCC model was established and mice treated with Enz (ENZ) or Olaparib (OLA) alone and combination. Bioluminescent images record tumor growth in different groups (three mice in each group). (B) Tumor growth was analyzed by relative emission of photons (left). Synergy analyses using Bliss independence model and two-way ANOVA analysis (right). (C) Immunohistochemcal staining for BRCA1 (upper), Ki67 (middle) and cleaved caspase-3 (lower), quantitations on the right. Quantitation data are presented as mean ± SD. *P < 0.05.

After sacrificing mice, we collected the orthotopic tumor samples to detect BRCA1, Ki67 and cleaved Caspase-3 in each group using immunostaining. BRCA1 staining in Enz alone group and Enz plus Olaparib group were significantly reduced compared with vehicle group **(Fig. 6C above)**which is consistent with our *in vitro* result. Ki67 level in Enz plus Olaparib group was significantly less than other groups **(Fig. 6C middle).**Cleaved Caspase-3 abundance in Enz plus Olaparib group was more than other group, although not reaching statistical significance **(Fig. 6C below)**. This induction of apoptosis likely was responsible for the tumor shrinkage, which could also be a consequence of direct anti-proliferation effect of the combination therapy.

Together, results from the *in vivo* animal model demonstrated that Enz plus Olaparib could lead to better suppression of the HCC growth, although a larger sample size will be needed to reach statistical significance for the synergistic therapeutic effect *in vivo*.

## Discussion

Most patients unfortunately receive a diagnosis of HCC when it has already reached an intermediate or advanced stage as HCC presents few symptoms in an early stage. Advanced HCC, due to limited treatment options, has very poor prognosis. Sorafenib (Nexavar), a multi-targeted kinase inhibitor, is the only drug approved by the FDA to improve outcomes in patients with advanced HCC^28^. However its effect with median survival extending only about 3 months is much to be desired.^29^, therefore, there is still a challenge for the development of new therapies for HCC.

Olaparib was approved for patients with BRCA-mutated advanced ovarian cancer and was also shown to be effective in BRCA-loss breast and prostate cancer^8,9^. However, its efficacy in HCC appears to be minimal due to a rarity of BRCA mutation in HCC based on deep sequencing^30–32^. In this study, we found that Olaparib could be selectively against AR-negative HCC cells, which might be due to AR’s regulation of BRCA1 and HR activity **(Fig. 1–2)**, similar to AR’s role in maintaining HR gene expression and activity in prostate cancer^33^. Indeed, a combination of AR and PARP inhibitor can synthetically kill castration-resistant prostate cancer^36^. These results suggest that Olaparib efficacy may not be strictly dependent upon a loss of BRCA1 gene, a lower BRCA1 expression might also benefit from a potential selective therapeutic effect of this drug, therefore identifying signaling pathways and compounds that can produce “BRCAness” will expand the use of Olaparib in other cancers through combination therapy. Alternatively, Olaparib single agent therapy in HCC will likely only benefit patients with AR-negative HCC, as previous studies showed that only about two thirds of HCC contained AR^34–36^.

Earlier studies demonstrated that the AR promotes HCC tumorigenesis and might be a potential therapeutic target for HCC^15,16^, but antiandrogen therapy failed in clinical studies^20–22^. In this study, we found that blocking AR signaling, although it could not dramatically induce HCC cell death **(Fig. 3B)**, likely inhibited BRCA1-dependent HR DNA repair **(Fig. 2)**. Using different HCC cell lines and approaches, we demonstrated that a new therapy combining the two FDA approved drugs, Enz and Olaparib, targeting different DDR pathways might be effective for advanced HCC **(Fig. 3)**. These findings may revive antiandrogens therapy as an adjuvant therapy for HCC. In addition, BRCA1 may not be the only factor that contributes to impaired HR caused by AR inhibition. Indeed, our data from TCGA database showed that in clinical samples AR was also associated with BRCA2 and RAD51 **(Fig. 2A)**. Therefore AR inhibitors combined with other mechanism-based DNA damage inducing treatment (chemotherapy drugs, radiotherapy) may also synergistically suppress HCC.

Various miRNAs are involved in HCC drug sensitivity. Our previous study demonstrated that miR-367 increased HCC Sorafenib chemotherapy efficacy^37^. In our study, we identified a new AR/miR146a-5p/BRCA1 pathway that contributes to the synthetic lethality of Enz and Olaparib. The miR-146a could down regulate BRCA1 in triple negative sporadic breast cancers^38^ and miR-146a-5p was also shown to be involved in liver inflammatory and HCC progression^39,40^. In **Fig. S1D**, we also showed that miR-146a-5p might be a protective factor for HCC development and progression. These data suggest that miR-146a-5p might be a crucial factor in HCC that should be considered as a potential target for HCC therapy. Indeed our work indicated that AR can regulate miR-146a-5p expression, thus clinically used drugs such as Enz can be applied to regulate expression of this miRNA to provide therapeutic gain for patients.

In summary, the findings presented here established that AR signaling could regulate BRCA1 and HR activity that might lead to differential sensitivity of HCC to Olaparib. Combining antiandrogen Enz with the PARP inhibitor Olaparib greatly enhanced the anti-tumoral activity in HCC. The mechanism study revealed that AR inhibition suppressed HR through the miR146a-5p/BRCA1 pathway **(Fig 7)**. Our preclinical data established a foundation for clinical studies with the combination of those two FDA approved drugs in HCC.

**Fig 7.**
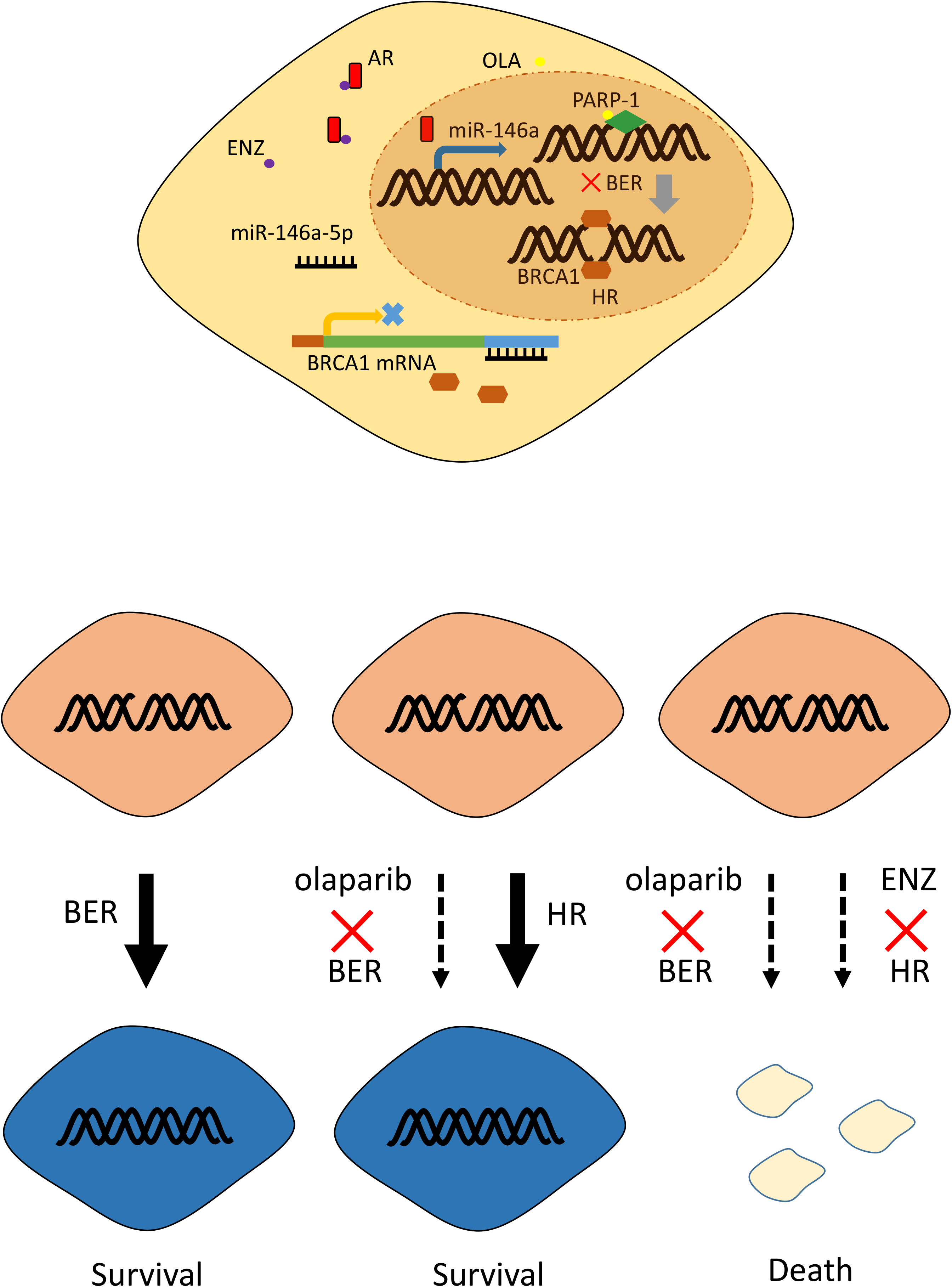
Model of Enzalutamide and Olaparib synergistically suppressing HCC. Enzalutamide (ENZ), as a selective antagonist of AR, inhibits AR binding with the promoter of miR-146a-5p and then increases its transcription. More miR-146a-5p targets the 3’UTR of BRCA1 mRNA. As a consequence, BRCA1 protein decreases. Loss of BRCA1 leads to HR deficiency. Olaparib targets PARP-1 leading to base excision repair (BER) deficiency. Combining ENZ with Olaparib will lead to irreparable DNA damage and cell death.

## Acknowledgments

This work was supported by NIH grant CA156700 and George Whipple Professorship Endowment, and Taiwan Department of Health Clinical Trial and Research Center of Excellence grant MOHW104-TDU-B-212-113002, and Zhejiang Provincial Natural Science Foundation of China LZ14H160002 and LQ18H160010.

---

**Fig S1.**
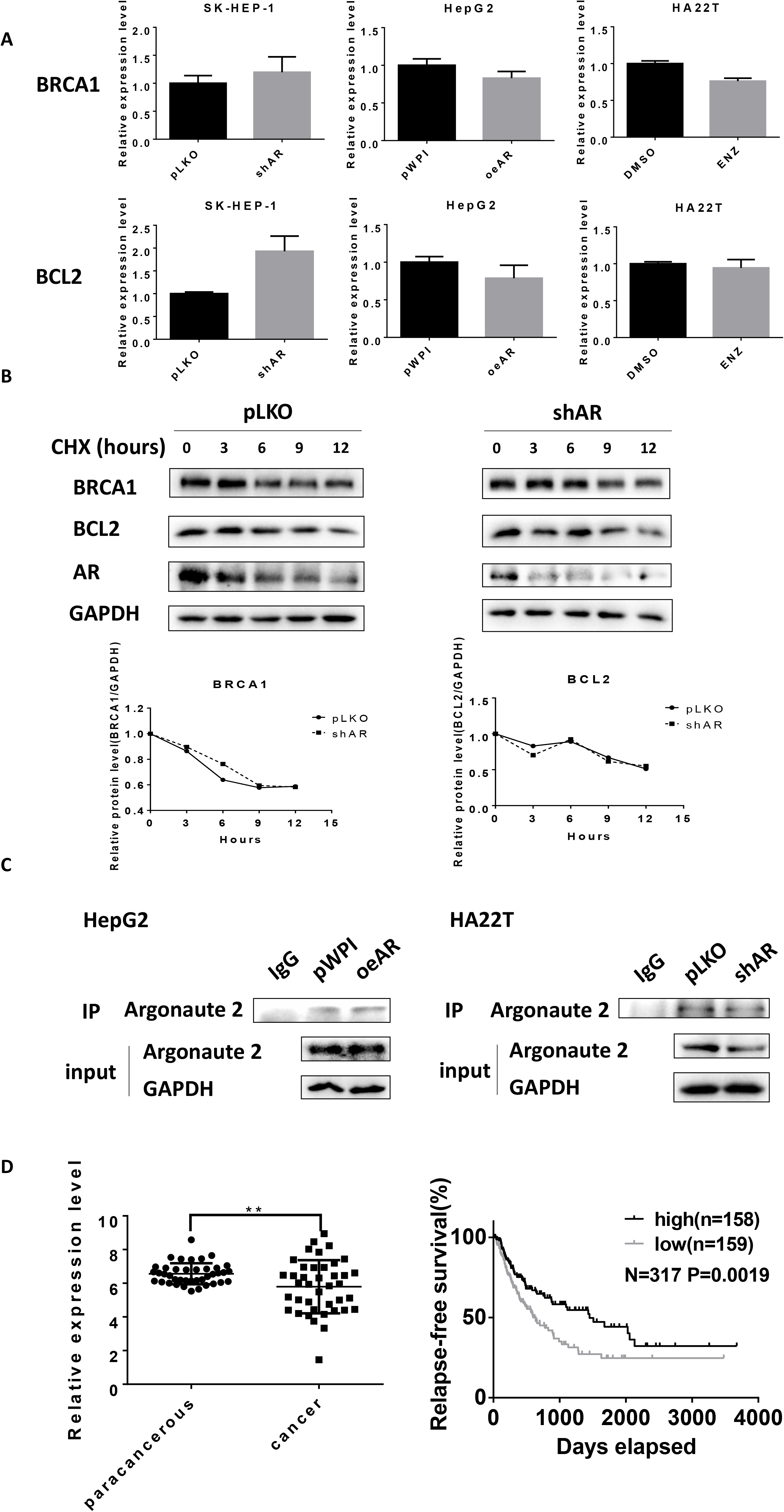
A) After manipulating AR or treating with Enz, mRNA levels of BRCA1 and BCL2 in SK-HEP-1, HepG2 and HA22T cells were detected using qRT-PCR. (B) Time course analysis of BRCA1 and BCL2 stability after cycloheximide (CHX) treatment in SK-HEP-1 cells. BRCA1, BCL2, AR and GAPDH protein levels at 0, 3, 6, 9 and 12 hours after CHX addition were analyzed by western blotting (upper) and quantified by ImageJ software (lower). (C) The protein level of Argonaute 2 in cells (input) and in pull-down beads (IP) during RIP assay was detected using western blots in HepG2 cells (left) and HA22T cells (right). (D) The miR-146a-5p expression level in HCC tumor and paracancerous tissues (left) from TCGA database, **P<0.01. The relapse-free survival of patients with high (the top 50% of total patients) or low (the bottom 50%) miR-146a-5p expression level (right).

